# Re-thinking recreational fishing – how a natural disaster presents insights and opportunities for achieving sustainability and equity objectives

**DOI:** 10.1101/2023.04.12.536519

**Authors:** Shane Orchard, Shawn Gerrity, David R. Schiel

**Affiliations:** University of Canterbury

**Keywords:** Coastal management, fisheries, public resources, open-access, beneficiaries, environmental change

## Abstract

Pāua (abalone) are a treasured natural resource that supports a recreational fishery worth $2 million annually to the local economy of the Kaikōura district in New Zealand. From 2016, the fishery was closed for 5 years in response to widespread mortality caused by co-seismic uplift in the 7.8 M_w_ Kaikōura earthquake. The fishery re-opened in 2021 for an initial 3-month open season with a recreational fishing allocation of 5 tonnes. We constructed scenario models informed by fishing pressure observations and show that this catch target was severely exceeded (by a factor of 9-10). We then evaluated a range of alternative management settings involving daily bag limit adjustments and temporal controls that shift opening times away from high visitation periods and favourable weather conditions as a means to reduce peaks in fishing pressure. Temporal control strategies reduced seasonal catch by up to 45% in comparative analyses using the 2021 fishing effort as a base-case scenario, and can be used in combination with daily bag limit adjustments and longer-term protected areas. When applied to a Total Allowable Catch, these solutions also produce equity effects that increase the proportional allocation accruing to local fishers. We discuss the potential role of these insights in designing temporal controls for managing seasonal influxes of fishing effort driven by tourism, and highlight the role of natural disasters and recovery periods as a trigger for new policy directions. In this case, they were needed to address an unexpected loss that illustrates a more general need for reliable fishery management that is responsive to environmental change in the long term.

## 1. Introduction

Open-access natural resources underpin community wellbeing and livelihoods throughout the world. They include a wide range of consumptive and non-consumptive resources often referred to as ‘commons’ (Folke 2007; Ostrom 1990). Despite being natural resources in public lands and waters, they are often the subject of controls on usage and exploitation to avoid ‘tragedy of the commons’ effects that arise when resource users lack incentives or requirements for ensuring the sustainability of such resources, or worse, engage in first-come-first-served extraction activities to the detriment of others (Hardin 1968). Across all public natural resources, recreational fisheries are among the most hotly contested due to the important role they play in many countries and communities (Barbier et al. 2011; Dyck & Sumaila 2010). Moreover, they are frequently the subject of long-standing traditional uses for indigenous peoples, often accompanied by traditional governance systems (Begossi 2014; McCarthy et al. 2014; Montgomery & Vaughan 2018; Raymond-Yakoubian et al. 2017).

In the face of an increasing human population and climate change, however, there are many challenges for natural resource governance (Hinkel et al. 2014). They include the need to balance increasing demands for use within a general context of environmental change and historical resource degradation. Many key aspects of these challenges have been explored and enunciated in a variety of seminal concepts that have helped to chart a way through these difficult waters. Examples include the concept of shifting baselines (Pauly 1995), merits of protected areas (MacKinnon et al. 2011; Watson et al. 2014), integration of protected areas and wider spatial planning (Douvere 2008), and necessity of recognising the interactions between social and ecological processes when devising resource management controls (McGinnis & Ostrom 2014). For the latter, the implementation step is invariably the locus of real progress. This is clearly seen, for example, in the functions of management agencies tasked with setting tangible controls such as access or use limits for fisheries (Beddington et al. 2005), and ideally evaluating their outcomes and adjusting as necessary in adaptive management cycles (Folke et al. 2005; Haasnoot et al. 2013). Unfortunately, there are many potential pitfalls in delivering these functions effectively. They include accounting for imperfect knowledge of the resource to be managed and balancing the competing demands of stakeholders while also remaining open to new ways of thinking about the management challenge and expanding the scope of the available management tools (Garcia & Charles 2008; Pita et al. 2019). Here, we address the latter in the context of the recreational pāua / abalone fishery (*Haliotis* spp.) along the Kaikōura coast in New Zealand.

The term ‘recreational’ fishery refers to the non-commercial harvest of fisheries that are available to the general public. Worldwide, there are an estimated 220 million recreational fishers or 5 times the number of commercial capture fishers (Arlinghaus et al. 2019; Arlinghaus et al. 2015). These fisheries are important at many scales and involve a wide range of coastal invertebrates and fishes (Arlinghaus et al. 2013; Ihde et al. 2011; Radford et al. 2018). In this case, recreational fishing is a prominent driver of tourism to the Kaikōura coast and this contributes to the variability in fishing pressure. Uniquely, the pāua fishery was closed for a 5-year period to facilitate recovery from the 7.8M_w_ Kaikōura earthquake which devastated the nearshore environment due to a combination of co-seismic sea-level changes and longer term erosion and sedimentation effects that had severe ecological impacts on habitat-forming seaweeds and other marine life (Orchard et al. 2021; Schiel et al. 2019; Schiel et al. 2021; Thomsen et al. 2021).

*Haliotis iris* (blackfoot) and *Haliotis australis* (yellow foot) are grouped together in quota allocations in the New Zealand pāua fishery. However, the vast majority of the fishery is for *H. iris*, as *H. australis* tends to occur as isolated individuals whereas *H. iris* can form dense aggregations. A novelty of *H. iris* is that it recruits from the plankton in very shallow water, and occupies specific under-boulder habitat from around the low tide mark to usually no more than about 3m depth (Gerrity et al. 2020). Juveniles live in these under-boulder habitats for about 3 years, before emerging at around 80-90 mm shell length, when they become reproductively mature (Wilson & Schiel 1995). These then meander along and down rocky reefs and can be found to 20 m depth in some locales (Schiel et al. 1995). However, much of the commercial fishery is in the 5-10 m depth range where most pāua aggregate. One of the unique effects of coastal uplift from the Kaikōura earthquake was the abundance of blackfoot pāua in shallow water in the recovering intertidal zone (Falconer et al. 2022; Gerrity & Schiel 2022). The closure of the commercial and recreational fishery took almost complete fishing pressure off pāua along the earthquake-affected coastline and similar abundances had not been seen since the early 1970s. Prospects for re-opening the fishery included the potential use of innovative management that may differ from the pre-disaster *status quo*. However, evidence for a substantially recovered pāua population perhaps underlaid the management approach for an initial 3-month re-opening of the recreational fishery with relatively few enforceable restrictions other than a slightly reduced daily bag limit in comparison to the pre-earthquake fishing rules. The associated outcomes and management implications provide the context for this study.

The objectives for our study were therefore to: a) assess fishing pressure across the entire management unit for open season, b) estimate the seasonal catch for comparison with the catch target, and c) evaluate alternative management approaches for their potential contribution to sustainable fisheries management with a focus on effective and equitable solutions for the achievement of catch targets. We conclude by discussing the most promising strategies for transforming the management setting towards a quantifiable, enforceable and ultimately more sustainable fishery.

## 2. Methods

### 2.1 Study area and context

#### Fishery management

The New Zealand pāua fishery is managed by the Ministry of Primary Industries (MPI) using a suite of controls that can vary according to the status of stocks within regions of coastline (Schiel 1992). The fishery is restricted to hand-gathering and snorkelling. There is a Total Allowable Commercial Catch (TACC) in each region annually, which is subject to adjustment depending on the status of stocks, and this is part of the Quota Management System. Commercial quota can be bought, sold and traded. There is a minimum legal size, which is generally 125 mm shell length (SL), although the commercial sector voluntarily raised this to 135 mm SL in the Kaikōura region post-earthquake to add further stock protection. There is also a customary fishing allocation to Māori, which varies in different stocks around the country. To promote recovery from the Kaikōura earthquake, the fishery was closed under section 11 of the Fisheries Act 1996 which contains provisions for sustainability measures (New Zealand Government 2022). The area subject to the closure involved c.135 km of coast within two fishery management units, PAU3A (the focus of this study), and a portion of PAU7 to the north (Figure 1).

**Figure 1.**
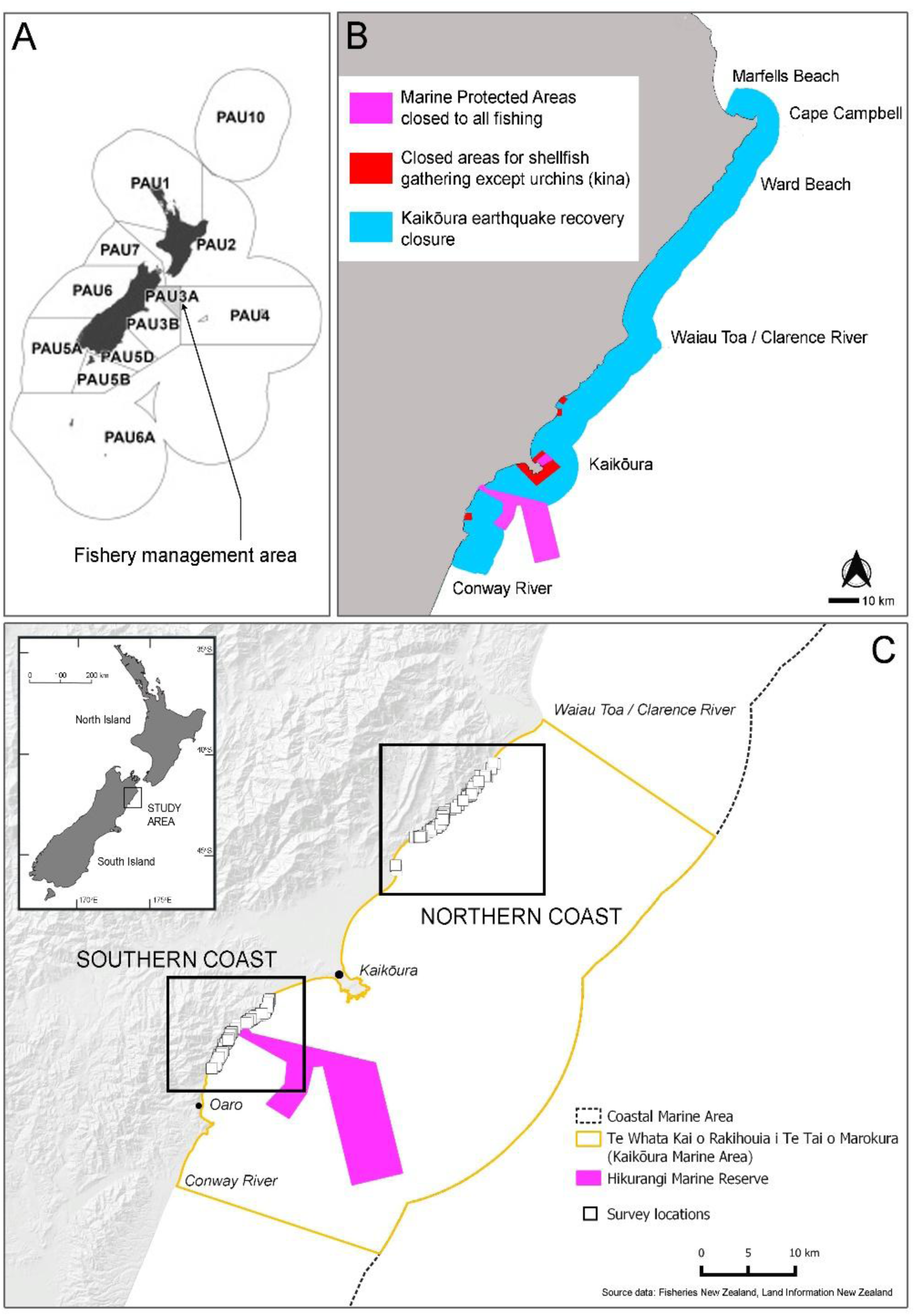
Maps showing the fishery management areas for pāua (A), the area closed to pāua fishing after the 2016 Kaikōura earthquake (B), and the survey locations for this study (C).

The fishery was initially reopened for the 3-month period of 1 December 2021 to 28 February 2022 (Figure 2). For PAU3A, a recreational fishing allowance of 5 t contributed to a Total Allowable Catch (TAC) of 40.5 t that included commercial (23 t), customary (7.5 t) and natural mortality (5 t) allowances. Recreational fishery rules included a minimum harvesting size of 125 mm shell length (SL), and a daily bag limit of 5 pāua per individual fisher. However, one individual could fish for many others so long as those people were present on the shore or in a boat. There was an ‘accumulation limit’ of 10 pāua, but this provision is almost entirely unenforceable. As is the case nationwide, none of the recreational catch was reportable and there is no marine fishing license requirement for individual fishers.

**Figure 2.**
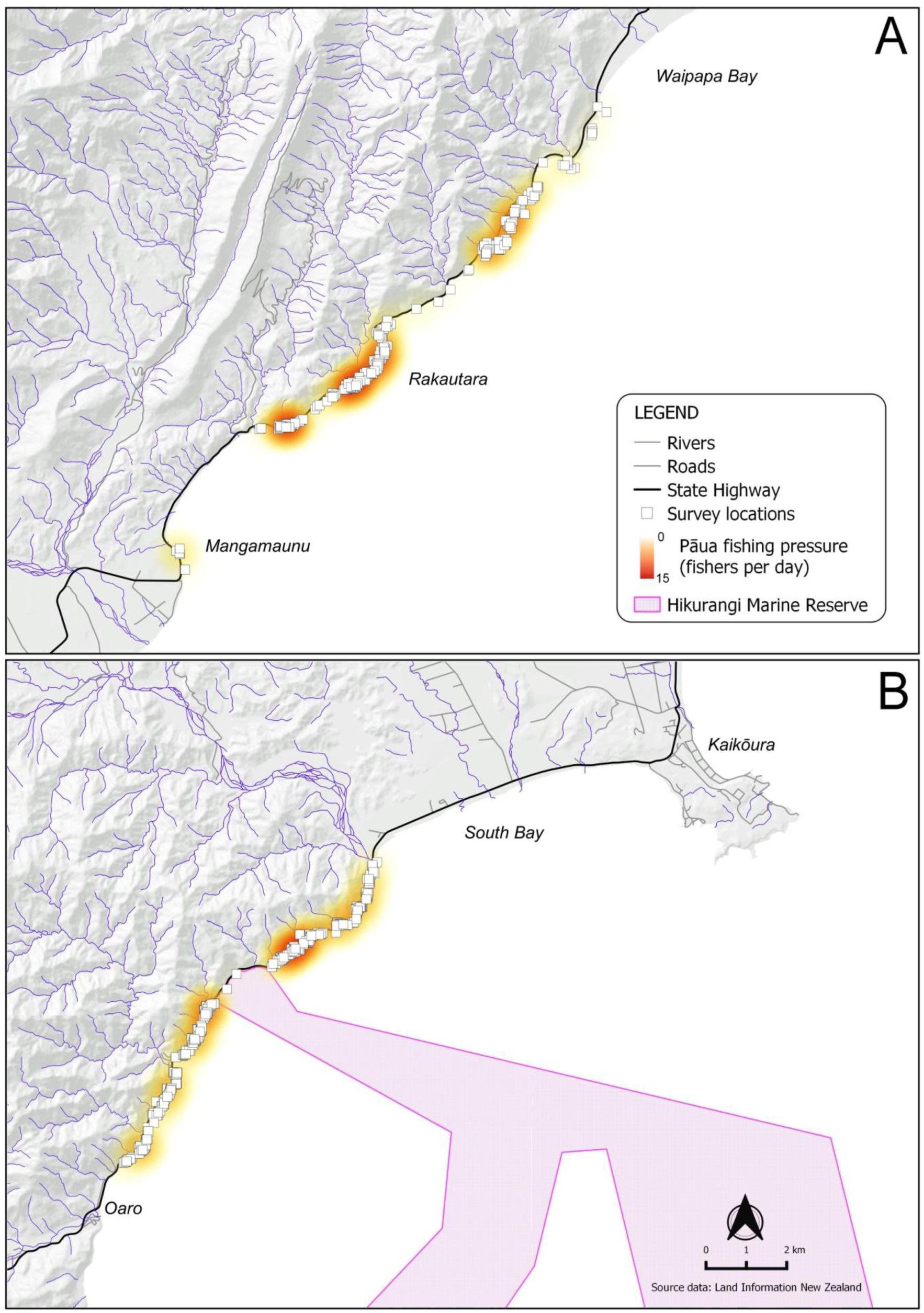
Survey areas on the northern coast (A), and southern coast (B), showing position of major roads and recreational pāua fishing pressure observed in the 1 December 2021 – 28 February 2022 fishing season. Note that pāua habitat on Kaikōura Peninsula (which is traditionally a pāua gathering area) remained closed to fishing during this period.

#### Study area

The study area was defined as the rocky reef and boulder habitats where recreational pāua fishing is done in / Kaikōura Marine Area (Te Whata Kai o Rakihouia i Te Tai o Marokura) between Waipapa Bay in the north and Oaro Beach in the south, with the exception of pāua habitat on Kaikōura Peninsula which remained closed to fishing (Figure 3). The study area represents practically all of the pāua habitat that was open to recreational fishing in PAU3A in the 2021-2022 season. To the north of Waipapa Bay there is a long stretch of predominantly mixed-sand gravel beaches (i.e., unsuitable habitat), that continue to the boundary with PAU7 (Figure 3A). To the south of Oaro Beach there are a few rocky reefs accessible from the public road but these fall within a mataitai (customary marine protected area) and were not included in the study area as they are subject to different fishery controls. The coastline further south is generally remote from the road and expected to receive considerably less recreational fishing pressure (Figure 3B).

**Figure 3.**
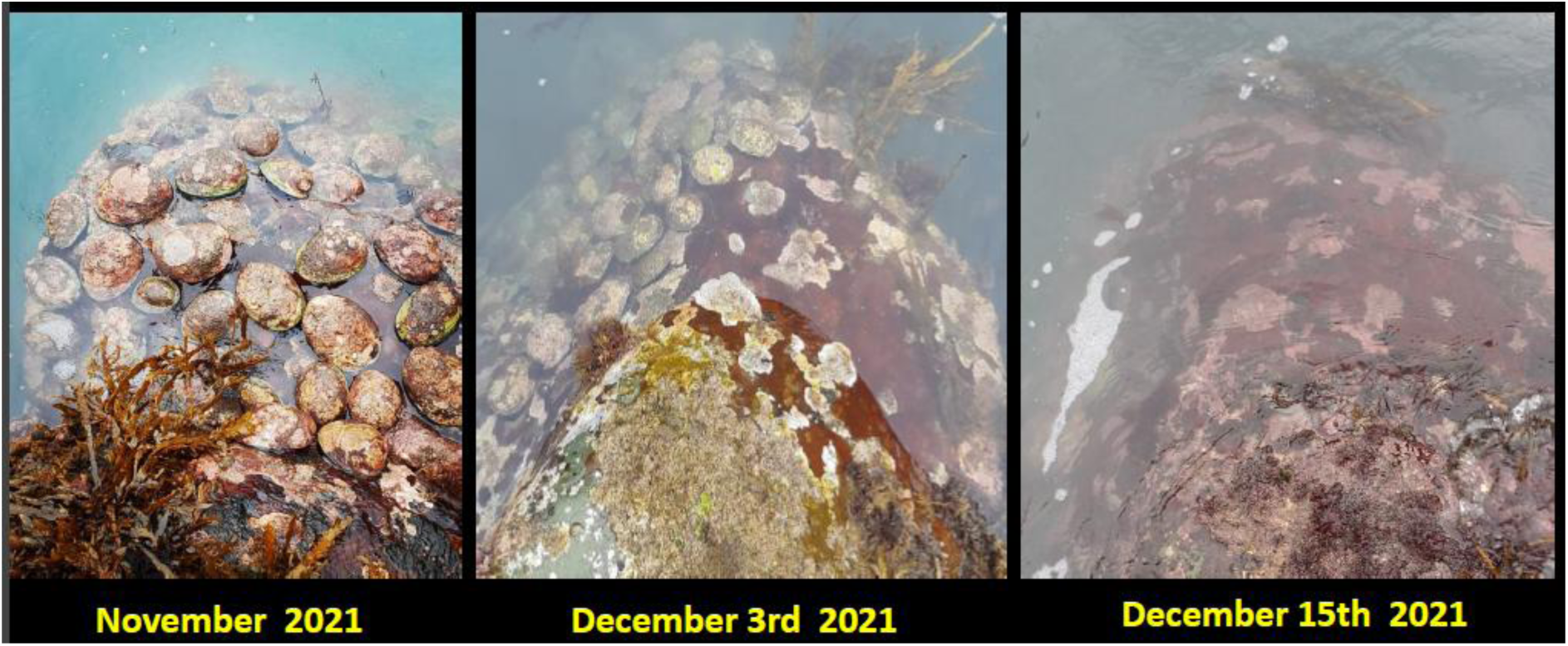
An example of the high pāua (New Zealand abalone) abundances that resulted from closure of the fishery for 5 years in response to the Kaikōura earthquake, and the speed at which harvest-size individuals were depleted following re-opening of the fishery on 1 December 2022. The abundance of accessible pāua was not the root cause of the overfishing that eventuated - rather, it was the lack of effective management for the level of fishing effort.

### 2.2 Fishing pressure assessment

Throughout the study area, all of the fishing activity was associated with vehicle access points on State Highway 1 (a distance of 56 km by road) which runs close to coast (Figure 3). Fishing pressure was estimated using surveys of fisher numbers across the entire study area on 19 days during the open season using a random stratified design that considered three visitation periods characterised as weekday, weekend and holiday. Within the 90-day fishing season there were 49 weekdays, 18 weekend and 23 holiday days based on a summer holiday period of 25 December 2021 – 16 January 2022, which reflects common workplace closing and re-opening times over the Christmas (summer) holiday period (Supplementary Material Table S1). Fisher numbers were recorded on the basis of short-duration ‘snapshot’ observation periods of around 5 minutes, each performed by a team of two people from vantage points at individual reefs along the coast. Each daily survey commenced at approximately 1.5 hours before the time of low tide from a random starting point and typically took 3-4 hours depending on the number of fishers present on the day. Each survey identified fishers engaged in pāua gathering, or gearing up or down with equipment for doing so. Other observed activities that were not identified as pāua fishing included spearfishing, line fishing or picnicking on beaches adjacent to areas of reef. Each pāua fisher was associated with a fishing group. These were identifiable on both land and in the sea from grouping and interaction clues, often assisted by the pattern of vehicles at road access points.

**Table 1.** Current management methods for the recreational pāua fishery in New Zealand. Source: Fisheries New Zealand.

| Fishery | Blackfoot pāua | Yellowfoot pāua | Details |
| --- | --- | --- | --- |

| management methods | (*Haliotis iris*) | (*Haliotis australis*) |  |
| --- | --- | --- | --- |
| Minimum size limit | 85 mm - Taranaki Region<br>125 mm - rest of New Zealand | 80 mm (all areas) | Applies in all areas not subject to no-take provisions (e.g., marine reserves) |
| Maximum catch per person per day | 10 | 10 | Applies in most areas<sup>†</sup> not subject to no-take provisions (e.g., marine reserves) |
| Accumulation (bag) limit | 20 pāua or 2.5 kg of shucked pāua (shell removed) | 20 pāua or 2.5 kg of shucked pāua (shell removed) | Applies to both species combined |
| Fishing method controls | Underwater breathing apparatus (UBA) cannot be used | Underwater breathing apparatus (UBA) cannot be used | UBA cannot be used while collecting pāua or while in possession of pāua. This includes strict controls on having a UBA in a vehicle or vessel that has been used for transport to a pāua fishing ground |

<sup>†</sup> with the notable exception of the area affected by the 2016 Kaikōura earthquake, including the Kaikōura (Te Tai o Marokura) Marine Area, as detailed in this study

To estimate the proportion of daily fishing effort represented in the snapshot surveys, continuous observations of fishing effort were made near Rakautara on the northern coast over the period 8 December 2022 – 9 January 2022 (Figure 2). These assessments used a combination of repeated site visits and timelapse photography from distant vantage points following New Zealand Privacy Commission and Department of Conservation guidelines. Fisher numbers, group size, arrival and departure times were tabulated to calculate the daily fishing effort and mean duration of fishing trips for comparison to snapshot survey data. From these data a daily visitation coefficient (VC) was developed to reflect the unobserved fishing effort per day within each visitation period. Values of 1.5, 2.0 and 2.5 were adopted as the best estimate of the mean VC for the visitation periods of weekday, weekend and holiday, respectively. These values were used to scale the observed fisher numbers within each visitation period to construct a seasonal fishing pressure model, and sensitivity analyses were then applied to explore the uncertainty contributed by variation in the assumed VC values (Supplementary Material Table S2). To account for the potential influence of inclement weather on fisher numbers, a total of six days were also deducted from the fishing pressure model based on an analysis of daily weather records from the Kaikōura Aws station (Lat -42.42038, Lon 173.69627) as recorded in the National Climate Database (https://cliflo.niwa.co.nz/). Bad weather days were identified on the basis of a minimum temperature <11 ^°^C coinciding with a maximum temperature <18 ^°^C (indicative of a cold day), or daily rainfall exceeding 20 mm (Table S3). For these days the daily fishing effort was assumed to be zero. This represents a conservative approach since some inclement weather days (e.g., one of rain and two of drizzle) were also among those sampled in the field surveys (Supplementary Material Table S1).

**Table 2.** Alternative management scenarios for the recreational pāua fishery on the Kaikōura coast.

| Management tool | Number of scenarios | Description |
| --- | --- | --- |
| Catch per day (CPD) limit | 6 | Daily limit ranging from 1 to 6 pāua per person per day |
| Season length | 4 | Season lengths of 9 – 12 weeks while retaining a 1 December start date<sup>†</sup> |
| No Fishing Days (NFDs) | 4 | 1 or 2 NFDs per week reflecting different weekend/weekday combinations |
| Timing of season | 2 | Timed to avoid or include the summer holiday period for season lengths of 9-13 weeks |
| Combinations | 174 | Combinations of the above |

<sup>†</sup> These scenarios reflect bringing forward the closing date by 1 – 4 weeks relative to the 2021-2022 season which ended on 28 February 2022.

**Table 3.** Estimates of seasonal catch patterns for the 2021-2022 pāua fishing season on the Kaikōura coast calculated from a fishing pressure model and incorporating three visitation periods and six catch-per-day (CPD) scenarios ranging from 1 to 6 pāua per-person-per-day.

| Tonnes | Number<br>of days | Catch-(tonnes) |  |  |  |  |  |
| --- | --- | --- | --- | --- | --- | --- | --- |
|  |  | CPD1 | CPD2 | CPD3 | CPD4 | CPD5 | CPD6 |
| Period |  |  |  |  |  |  |  |
| December | 31 | 2.8 | 5.6 | 8.4 | 11.2 | 14.1 | 16.9 |
| January | 31 | 4.7 | 9.3 | 14.0 | 18.6 | 23.3 | 27.9 |
| February | 28 | 1.5 | 3.1 | 4.6 | 6.1 | 7.6 | 9.2 |
| Total | 90 | 9.0 | 18.0 | 27.0 | 36.0 | 45.0 | 54.0 |

(b) Catch by visitation period
| Tonnes | Number<br>of days | Catch-(tonnes) |  |  |  |  |  |
| --- | --- | --- | --- | --- | --- | --- | --- |
|  |  | CPD1 | CPD2 | CPD3 | CPD4 | CPD5 | CPD6 |
| Period |  |  |  |  |  |  |  |
| Holiday | 23 | 5.5 | 10.9 | 16.4 | 21.8 | 27.3 | 32.7 |
| Weekend | 18 | 1.9 | 3.9 | 5.8 | 7.7 | 9.7 | 11.6 |
| Weekday | 49 | 1.6 | 3.2 | 4.8 | 6.4 | 8.0 | 9.6 |
| Total | 90 | 9.0 | 18.0 | 27.0 | 36.0 | 45.0 | 54.0 |

### 2.3 Scenario analyses

Four sets of scenarios were developed to investigate the potential contribution of fishery management innovations for the general objective of reducing fishing pressure to achieve the Total Allowable Catch (TAC). The primary touchstone for interpreting the scenarios involves comparisons with the base-case. In this regard, the establishment of a time-bound open season with an associated catch target provides a unique opportunity for assessing the relative effectiveness of various management approaches in this fishery, since it is controlled in an open-ended manner elsewhere in the country.

#### Base-case scenario

The base-case scenario represents the estimated catch using the above fishing pressure model and a catch-per-day (CPD) of 5 pāua per-person. This reflects a scenario where the CPD is fully allocated as per each fisher’s personal entitlement and is likely to closely approximate the actual take in this particular case because of the abundance of easily captured pāua in shallow water. In addition, the adjustment of CPD is one of the key management alternatives to be considered and the base-case scenario is designed to facilitate this assessment. Catch estimates from the base-case (and other) scenarios were converted to biomass using a rounded figure of 3 pāua / kg. This represents a mean length-weight relationship of around 132 mm shell length following Schiel & Breen (1991) and is considered to be a suitable estimate of the average (legal) catch in these scenarios. This is slightly lower than the average shell length (138 mm) measured in catch sampling during the 2021-2022 season (Holdsworth 2022), which reflects the abundance of relatively large pāua and implies a higher catch weight. The figure we have adopted is more consistent with past catches and is more appropriate for the forward-looking scenarios presented here.

#### Alternative management scenarios

Limits on the CPD allowance provide one of the most obvious and traditional management tools for the control of fish pressure. In this case, a small adjustment to the CPD limit was one of the primary management approaches used in the strategy for re-opening of the fishery, in addition to the limited open season. It involved a modest reduction to 5 pāua from the previous limit of 6 pāua that had been in place within the Kaikōura Marine Area since 2014 (New Zealand Government 2021), and this was already more stringent than the national regulations in force elsewhere (10 pāua per-person-per-day, Table 1). All of our scenarios involved interactions between various temporal controls and six CPD scenarios (daily limit 1 to 6 pāua per-person). Four shortened-season scenarios were considered that involve reducing the season length by bringing forward the closing date while retaining a 1 December opening date as used in 2021. These reflect season lengths of 9-12 weeks relative to the base-case of 13 weeks (Table 2). The scenarios for season-shortening were not extended into the holiday period, but other options for reducing fishing pressure in the holiday season were considered including the use of No Fishing Days (NFDs) and alternative seasonal timing. We considered four NFD settings that reflect the establishment of either one or two NFDs per week for the duration of the season, representing different weekend/weekday combinations, and interactions between the NFD settings, CPD and season length. Finally, the effect of moving the season to outside of the holiday period was explored with season lengths of 9-13 weeks, along with various combinations of the CPD and temporal controls in place (Table 2).

## 3. Results

### 3.1 Fishing pressure

#### Coast-wide survey results

Fisher numbers observed during the coastwide snapshot surveys ranged from a high of 497 on 2 January 2022 to a low of 11 on 27 January 2022. Mean values (±SE) for the three visitation periods were 73 ± 12 (weekday), 174 ± 76 (weekend) and 285 ± 79 (holiday). The evidence collected from repeat surveys and timelapse observations suggested that the daily visitation pattern typically involved around twice as many fishers as was observed during the snapshot surveys, and often more on busy days. At Rakautara, a relatively popular location with abundant pāua, a consideration of arrival and departure times showed only a loose alignment with the time of low tide, with the overall pattern being consistent with an elevated period of fishing effort on the lower half of the tide and occasional arrivals at other times (Figure 4). The mean duration of groups on the reef was 42.5 minutes (±3.3 SE) (Figure 4B). However, since each individual group also spent time gearing up and down after arrival at the site, it was assumed that they were present on-site for an average of twice this length of time (90 minutes), during which time they could be observed in the coastwide snapshot surveys. This figure was compared with the cumulative fishing time recorded for all groups per day which averaged 144 minutes ± 44 SE for weekdays, 225 ± 130 for weekends and 258 ± 50 for the holiday period. The estimated VCs based on these calculations are 1.6, 2.5 and 2.9 for weekdays, weekends and holidays, respectively, at Rakautara. In our best estimate for the whole coast, slightly lower VCs (1.5, 2.0 and 2.5) were adopted to better reflect the contribution of less popular fishing locations. These VCs represent mean values across all sites and for which uncertainties were further investigated using sensitivity analyses (Supplementary Material Table S2).

**Figure 4.**
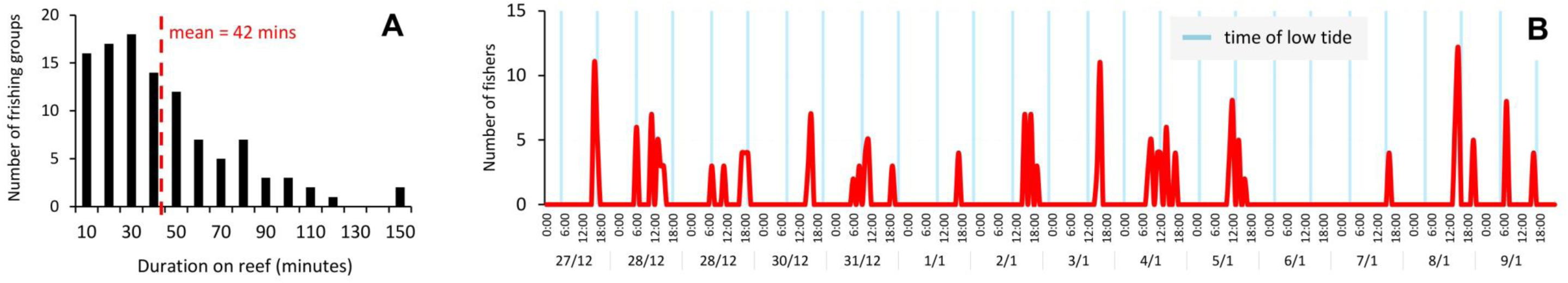
Temporal patterns in daily fishing activity from continuous monitoring of a reef near Rakautara. A) Histogram of the time spent by individual pāua fishing groups. B) Arrival times of fishing groups over the peak holiday period (27 December 2021 – 9 January 2022) in relation to the time of low tide (shown by the blue bars).

### 3.2 Seasonal catch model

#### Fishing pressure

The fishing pressure model reflects the estimated daily fishing pressure based on field survey and visitation coefficient calculations (Figure 5). At the opening of the season the total fishing pressure across the coast was modelled at ∼100 fishers-per-day with modest increases on weekends and good weather days. In the week before Christmas numbers started increasing and are initially under-predicted in the model until the modelled holiday period begins, starting on Christmas Day. Field observations showed huge pressure on the following days with numbers peaking at over1000 fishers on the busiest days and staying relatively high for most of January due to settled weather conditions. Our estimate for the modelled holiday period through to 16 January was 712 fishers-per-day. These estimates correlate well with field observations made by Fisheries Officers and numerous media articles that reported observations of >1000 fishers-per-day. The ‘weekend’ visitation period in this model showed the most variability with a mean of 341 fishers-per-day ± 155 SE (Supplementary Material Table S3). This was influenced by a noticeable peak in late February which coincided with good weather and may have been affected by the prospect of the upcoming season’s end (Figure 5).

**Figure 5.**
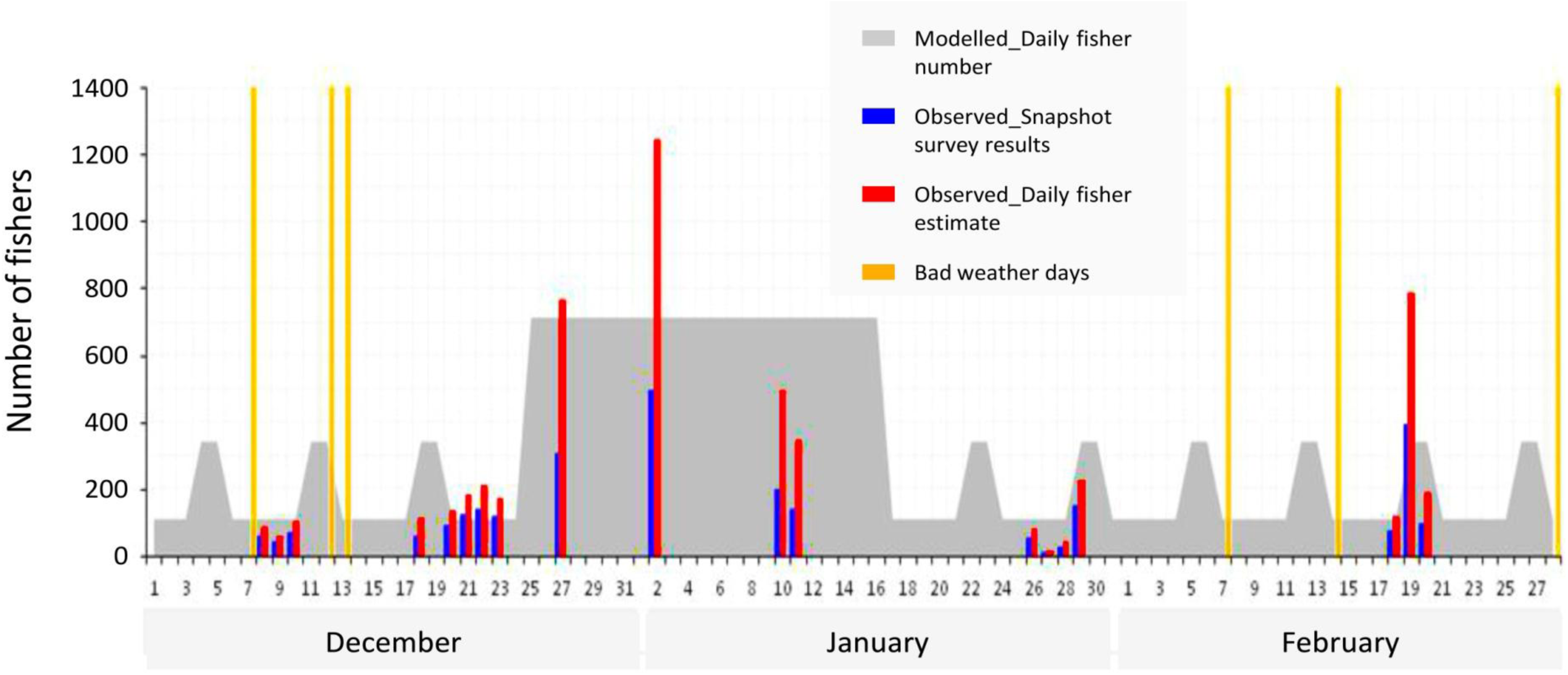
Fishing pressure model for the recreational pāua fishery during the 2021-2022 open season on the Kaikōura coast.

#### Base-case catch model

The base-case catch model assumes that the resource is fully utilised to the CPD limit of 5 pāua per-person (CPD5 scenario, Figure 6). Using the seasonal catch estimate of ∼135,000 pāua, this results in total catch weight of 45 t. Around half of this total was harvested during the month of January, followed by December (31% of seasonal catch) and February (17%), according to this model (Table 3). If the CPD is taken to be 4.5 pāua per-person as estimated from a limited catch sampling programme by Holdsworth (2022), then our seasonal catch estimate is 40.5 t. This is slightly lower than the 42 t seasonal catch estimate made by Holdsworth (2022) and reflects the slightly higher average weight of pāua in their model (as above) and sampling of elevated fishing pressure in the first week of the season which we did not specifically include. Nonetheless, the very similar results obtained from the two independent catch estimation programmes provide a high level of confidence for interpreting our base-case catch model and comparative scenarios. Potential uncertainties in our seasonal catch estimate are considered to be in the order of ±25% in consideration of variation within each temporal stratum and the upper and lower bound estimates for the visitation coefficient (Supplementary Material Table S2).

**Figure 6.**
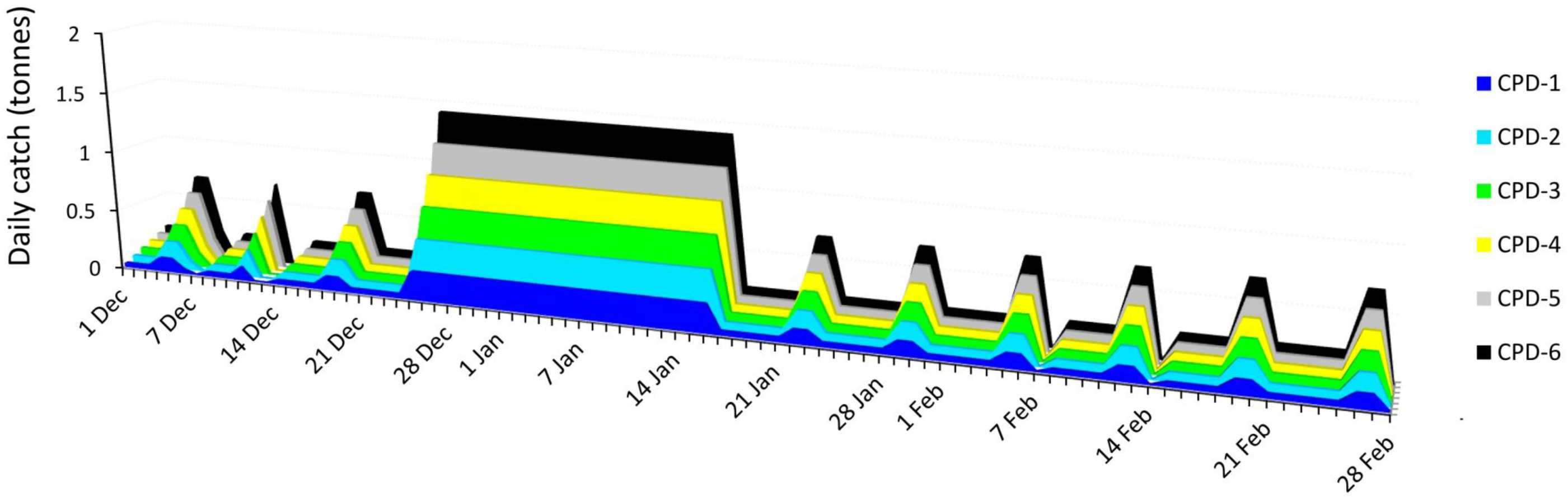
Seasonal catch model for the recreational pāua fishery on the Kaikōura coast based on fishing pressure observed during the 2021-2022 open season and six catch-per-day (CPD) scenarios ranging from 1 to 6 pāua per-person-per-day.

### 3.3 Scenario analyses

#### Daily bag limit adjustments

Reducing the daily bag limit is the most obvious way to reduce the total seasonal catch provided that it can be effectively implemented. The scenario analyses show that severe reductions would be needed to reduce the total catch to the target level of 5 t since even the most stringent (CPD1, an 80% reduction) generates a seasonal catch of 9 t (Table 3). This illustrates the need for other management innovations in addition to daily bag limit adjustments, such as the temporal controls we explore further below.

#### Season length and No Fishing Days (NFD)

Reducing the season length by bringing forward the closing date could produce modest catch reductions of 0.4 – 1.5 t for season length reductions of 1 - 4 weeks, respectively (Figure 7a, Supplementary Material, Table S4). For example, the shortest (9-week) scenario, with a closing date of 30 January, resulted in an estimated seasonal catch of 37.3 t representing a 17% reduction. Naturally, a much greater proportional reduction could be achieved if the shorter season extended across the busy holiday months when fishing pressure was higher, and this effect is partly seen in the NFD results. As an alternative to shortening the season, NFDs can be applied to reduce or maintain the number of available fishing days over a longer overall season on an annual basis. Scenarios we tested include a Sunday+Monday closure for the duration of the 2021-2022 season that resulted in a 40% reduction in seasonal catch assuming the is no compensatory shifting of effort (Figure 7b). The NFD scenarios also produced examples of a seasonal catches nearing the management target of 5 t. However, a combination of two NFDs per week and the most stringent CPD1 of 1 pāua per-person were needed to bring the estimated seasonal catch to under 6 t (Figure 7b).

**Figure 7.**
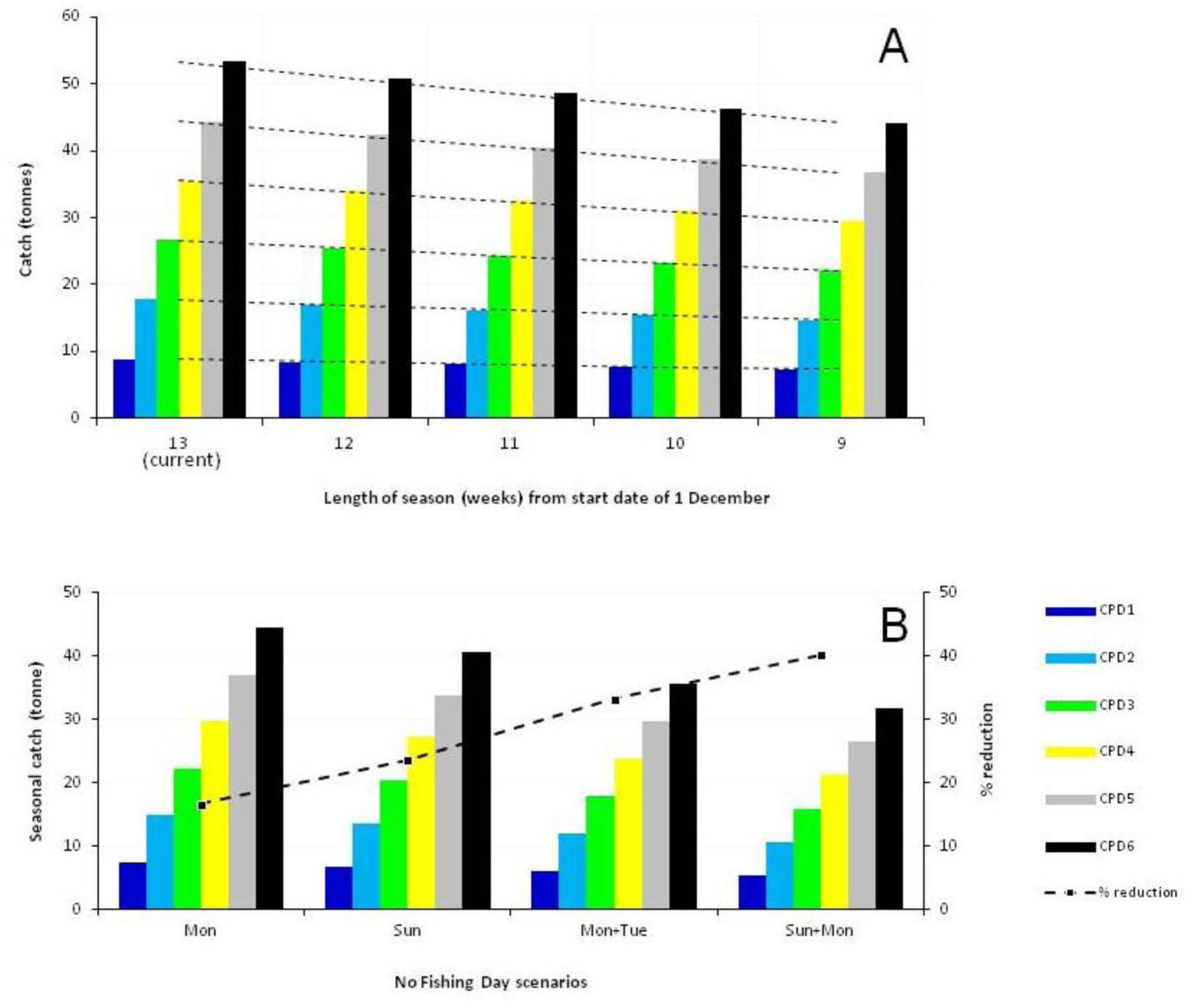
Examples of scenario analyses showing the effect of alternative management arrangements relative to those used in 2021-2022 recreational pāua fishing season on the Kaikōura coast. Both sets of analyses show the interaction with six catch-per-day (CPD) scenarios reflecting the daily bag limit in force. (a) No Fishing Day scenarios involving different weekend/weekday combinations. (b) Effect of shortening the season by bringing-forward the season closing date relative to the 2021-2022 fishing season which closed on 28 February 2022.

#### Seasonal timing scenarios

Similar to the effect of NFDs, moving the season to avoid the period of highest fishing pressure offers an effective route for reducing the seasonal catch and can be used in combination with the season length and daily bag limit adjustments. The 23-day holiday period accounts for 61% of the base-case scenario and the month of January accounts for 51% (Table 2). Using our catch estimate for February as an example of a non-holiday month the seasonal catch could be reduced by 45% by avoiding the holiday period while retaining a 13-week season length (Supplementary Material, Table S4). With this option the CPD1 scenario achieves the catch target of 5t while all shorter season length scenarios result in further catch reductions. This illustrates how a combination of seasonal timing and shorter season or NFD options could be used to achieving the seasonal allowance while providing for daily bag limits of >1 pāua per-person which is desirable in consideration of travel expenses and other costs associated with daily fishing.

## 4. Discussion

### 4.1 Interpretation of scenarios

For the purposes of this study based on comparative scenario analyses, an estimate of the 2021-2022 seasonal catch was used as a base-case to explore the relative merits of management alternatives. This back-casting approach carries the inherent assumption that future fishing pressure patterns would resemble the base-case (although others can also be considered), and also assumes that the core management objective of achieving control over the actual seasonal catch will remain relevant, which we believe will always be the case. Our seasonal catch estimate represents a defensible total despite the challenges inherent in obtaining an accurate measure due to variation in the timing and duration of daily fishing effort in relation to the short-duration ‘snapshot’ surveys upon which the field observations are based. As it is not logistically feasible to obtain a direct measure of coast-wide fishing pressure through field observations (i.e., requiring continuous monitoring of the whole coastline over the whole fishing season), this must be addressed by other means. These temporal aspects were addressed by developing a visitation coefficient that is consistent with time series data from a representative site and impressions gained from repeat field observations made at several sites on different times of the tide. These showed that the visitation pattern often involved the period of fishing effort commencing around mid-tide and continuing through the lower half of a tidal cycle. This approach facilitated the estimation of daily fishing effort while also identifying the major sources of uncertainty for extrapolation of the snapshot data to a daily time frame (Supplementary Material, Table S2).

A different set of assumptions is inherent in the three visitation periods used in the seasonal catch model. Their utility matches well with field observations made by our research team but potentially underestimates the length of the busy ‘holiday’ period. This is typically around 3 weeks (as assumed here) as a consequence of the timing of public holidays in New Zealand. In this case, the period of high visitor numbers and associated fishing pressure appeared to last an extra week before dropping off at the end of January – an effect that was likely influenced by a run of very good weather at that time. These impressions on the nature and timing of busy periods were also informed by discussions with MPI Fisheries officers who were working in the field performing compliance functions on all days of the fishing season. We did not specifically sample the ‘pulse’ of fishing effort that was expected in response to lifting of the fishery closure (i.e., early in the open season) but did complete field surveys towards the end of the season which showed evidence of elevated fishing pressure at that time.

### 4.2 Insights from scenarios

#### Effects of excessive pressure

Considering our seasonal catch estimate and potential uncertainties, the overriding conclusion from this study is that the recreational catch vastly exceeded its allowance and a similar estimate (42 t) was calculated independently by Holdsworth (2022). The excess fishing reflects the effects of unexpected and uncontrolled fishing pressure. At around 3 paua per kilogram, this represents well over 100,000 individuals, each of which takes around 7-8 years to reach the minimum harvestable size (Schiel 1992). It also represents an unplanned loss of spawning biomass that can have population feedback consequences leading to recruitment limitations and further decline (McShane 1992). Therefore, it is abundantly clear that alternative approaches are urgently needed to improve management. The scenario analyses presented here provide a methodology to identify lessons from the 2021-22 re-opening and promising alternatives.

#### Alternative management strategies

The single-most effective measure for achieving catch reductions relative to the base-case involves reducing the CPD. This is also a relatively equitable measure since it exerts a similar effect on all user groups across all days of the season. However, it is interesting to consider the potential contribution of a shorter fishing season, or adjustments to the seasonal timing, when used in combination with the CPD. In our scenarios, for example, just a single NFD per week has the potential to reduce seasonal catch by 16% for weekdays or 23% for weekends (Table 4). This strategy effectively reduces the number of fishing days within an open season while maintaining the opening and closing dates that represent a longer period. This could be seen as desirable in terms of spreading out both the fishing effort and the opportunity to go fishing over a longer time period. In contrast, reducing the number of fishing days by bringing forward the season closing date has the effect of compressing the fishing opportunity into a shorter period and was also less effective than NFDs for the objective of reducing seasonal catch. This result reflected the lower catch rates in February that are attributable to reduced fishing pressure following the peak summer holiday period. Since this general pattern can be expected in future years, the analyses suggest that temporal controls might offer significant benefits.

An extreme example of temporal controls has been used in the Roe’s abalone (*Haliotis roei*) recreational fishery near Perth in Western Australia. The area concerned was closed to both commercial and recreational fishing in response to severe mortality caused by a marine heat wave in the summer of 2011 (Hart et al. 2018; Strain et al. 2019). Although the recreational fishery has been re-opened, it is open for a specific number of 1-hour sessions on select days, requires a fishing licence and reporting, and is subject to a Total Allowable Recreational Catch (TARC) limit (Government of Western Australian Department of Fisheries 2021). The TARC also facilitates catch-sharing between the commercial and non-commercial sectors. As with New Zealand’s pāua fishery, the recreational catch proportion is substantial, with over 40% of the catch accruing to the recreational sector (Strain et al. 2021). The harvest strategy also recognises the limited recreational effort (15% of total) exerted in the subtidal fishery which supports 60% of the spawning biomass. The recreational allowance reflects a partitioning of effort between intertidal and subtidal areas for the benefit of recreational fishers who are afforded the easier access (Government of Western Australian Department of Fisheries 2021).

#### Social aspects and benefit-sharing

In the Kaikōura region, and for wider New Zealand, discrimination between intertidal and subtidal pāua stocks is not formally incorporated in fisheries management regulations but it is considered among the voluntary measures used by the commercial fishery to facilitate catch spreading. In cases of overfishing, however, the spatial pattern of impacts is important since these may exert even effects across the fishing community. An example from this case concerns the ability of less mobile (e.g., elderly) fishers who do not have the physical means or equipment to access subtidal stocks and are more reliant on harvesting from intertidal populations that offer a wadeable fishery. The prospect of maintaining a wadeable fishery may also support Māori whānau (families) who customarily use the shallow areas as a living larder for frequent harvesting, as with many other mahinga kai resources (McCarthy et al. 2014). These demographics contribute to the social impacts associated with overharvesting in specific areas, further emphasising the need for management strategies that can effectively achieve agreed targets for benefit-sharing.

Socio-cultural aspects of the fishery are further exemplified in our temporal analyses which highlight the relative fishing effort exerted (either intentionally or unintentionally) by various stakeholder groups, and opportunities to manage the benefits equitably. For example, a downside of the shorter season scenarios we tested involves disproportional effects on the local community members who may otherwise remain active in the late summer months and beyond. Conversely, shifting the season to outside of the holiday period would tend to support local community interests whilst reducing the allocation that is ultimately harvested by holiday-makers. These dimensions illustrate the importance of addressing fishing pressure and allocation demographics within the recreational sector that are additional to quota-management decisions between commercial and non-commercial sectors. Equity issues involving the distribution of benefits to participants are typical of public resources (Kockel et al. 2020; Yang et al. 2015), and may be particularly important in depleted or recovering fisheries where recreational and commercial sectors must work in concert to achieve sustainability objectives despite a high demand for access to the resource (Cooke & Cowx 2006; Ihde et al. 2011).

In seeking equitable benefit-sharing between commercial and non-commercial sectors, at least part of the challenge is dealing with the commonly-held and often expressed belief that recreational fishers have lower impacts on fish stocks and ecosystems (Arlinghaus et al. 2019; Cooke & Cowx 2006; Morales-Nin et al. 2015; Radford et al. 2018). However, this assumption cannot be tested unless it is actively measured. Stock assessments for New Zealand pāua fisheries generally use catch-per-unit-effort (CPUE) and length-frequency data from reported catches in the commercial fishery (Neubauer 2020), and with the non-commercial contributions estimated from the customary fishery records and representative studies (Fu et al. 2017). Recreational pressure and/or catch has been studied using creel surveys and catch diaries (Carbines 2000), or with off-site surveys (Heinemann et al. 2015), but may lack the spatio-temporal resolution needed for effective adaptive management. This case shows that extreme events and their recovery contexts may require additional survey effort to understand both the initial impacts and recovery dynamics in the affected social-ecological system, which can include influences from a wide surrounding region.

In this case, both commercial and non-commercial fishing sectors have an interest in recreational allowances that are established because of their contribution to the TAC for a given Fishery Management Area. It is also clear from case law that allowances set under the Fisheries Act 1996 are to be regarded as hard limits to be monitored and enforced where needed^1^. Therefore, the overfishing of an allowance by one sector has impacts on other stakeholders because it may reduce the TAC available in subsequent seasons. The challenges involve not only the proportional allocations to the commercial, cultural and recreational sectors, as is often argued, but how they can be managed as ‘one fishery’ where the management targets are achieved by all sectors. Additionally, quantification of the actual catch is also a key ingredient for sustaining fisheries in a changing environment. It facilitates, for example, timely responses to evidence for unexpected losses from either anthropogenic or naturally occurring stressors.

#### Interaction with protected areas

Protected areas are an important component of fisheries and wider ecosystems management, but may displace fishing effort into the remaining open areas (Abbott & Haynie 2012; Bastardie et al. 2014; Horta e Costa et al. 2013). Consequently, they may be best conceived as a supporting measure for a core set of fishery regulations that address the wider environment. Although spatial protections can help to increase fish stocks within this wider area through overcoming recruitment limitations, safeguarding nursery habitats or food sources, and the spillover of harvestable individuals (Gaines et al. 2010; Halpern et al. 2010; Jennings 2000), specific assessments are needed to determine the existence of these potential benefits (Bohnsack 1998). In this case, the relatively immobile nature of harvestable–size pāua populations suggests that spillover effects are unlikely to offset local depletion to a substantial degree, and that the benefits of long-term closures would relate mostly to recruitment effects from the protection of spawning biomass. In comparison, the temporal closures discussed here apply a consistent level of protection across the wider environment with the objective of preventing the serial depletion of spawning biomass in suitable habitats. Through a focus on fishing effort and pressure, these approaches can facilitate three of the four key strategies for conserving sedentary invertebrates identified by Jamieson (1993): high minimum size limits, unexploited refuges, maintenance of high densities in many areas, and the specific management of metapopulation units.

### 4.4 Adaptive and responsive management

Extraordinary circumstances offer new opportunities for innovation, and yet clear perceptions of the problems are a pre-requisite to formulating effective solutions. The initial 3-month opening of the New Zealand pāua fishery on the Kaikōura coast has provided a salutary lesson in what does not work well, and a cautionary tale of wishful thinking based on incorrect assumptions. While the methods and results of this initial post-earthquake fishery opening are clearly unsustainable they have helped to illuminate management shortcomings that may be important elsewhere. These include (i) reliance on the ‘accumulation limit’ concept which is essentially unenforceable except in overt circumstances, (ii) the need for effective ‘party’ limits to restrict a single person from gathering the bag limit for many other people in a group, (iii) the lack of a catch reporting facility, so that the extent of the fishery needed to be modelled from visitation and catch sampling assessments made largely after-the-fact, and (iv) the lack of a legislative mechanism to proactively close the fishery as needed to address higher-than-expected fishing pressure.

Disaster recovery and risk management aspects of this case include the restorative settings that initially led to resource recovery but were subsequently jeopardised in the transition to previous resource use levels. The statutory objectives for this recovery phase provided for resource use at 50% of pre-disaster levels within the context of sustainability measures (Fisheries New Zealand 2021). Key learnings which are likely applicable to other natural resource recovery situations, include the unreliability of daily allocations (e.g., bag limits) as the primary management tool in the absence of stable demand and pressure, which is perhaps more the rule than the exception in a changing environment. Additionally, New Zealand’s recreational pāua fishery is currently characterised by the lack of a mechanism to report catches or to allocate a recreational seasonal allowance on a per-fisher basis through strategies such as California’s tag-based system (Lewis 2015). It instead uses a daily bag limit for individual fishers as the key management strategy within an otherwise open-access fishery, and this carries the risk of overharvest if fishing pressure is greater than expected. If retaining this management approach, it would make sense to include a responsive policy provision that provides a regulatory backstop informed by catch reporting and/or catch sampling data, ideally in real-time, and could be triggered to adjust key management settings such as the season closing date, in response to evidence that the recreational allowance had been fished to its limit. Conversely, it could also be used to prolong the season should there be evidence of under-harvest due to factors such as inclement weather conditions. One of the key advantages of this approach is that allows for the selected management settings to be used in a more experimental manner, whereby learning is gained from their implementation, while also retaining control over the total seasonal catch, which is a priority for all stakeholders.

The traditional way of managing the NZ pāua recreational fishery through daily bag limits will most likely reduce the recreational catch, as shown here, but cannot completely address the problem unless it is set very low, which introduces inefficiencies for fishers in obtaining a ‘feed’, especially where they have travelled some distance to do so. Therein lies a conundrum since the ‘accumulation limit’ concept - which could in theory support an appropriate balance - is fundamentally unenforceable and therefore unreliable. At the same time there is an opportunity to trial new innovations such as the temporal closure strategies outlined here, and App-based reporting of catch and effort data during the season. In this way, the recreational fishery can be better understood and managed alongside the commercial and cultural components of the Total Allowable Catch for the season. This form of engagement with fishers may offer additional benefits for regulatory compliance by improving the awareness of individual impacts on the long-term future of fisheries, and allowing recreational fishers to have direct input into fishery management through providing critical data.

On a positive note, the Kaikōura coast example also shows that natural recovery can be generated through restorative management that controls anthropogenic pressures. This included the re-establishment of a wadeable fishery of the likes not seen for decades. In the case of pāua in New Zealand, the shallow water populations have mostly disappeared or become severely reduced in accessible places. The Kaikōura coast experience illustrates the potential for natural recovery to reverse historical decline and deliver abundances never before seen by the younger generation of fishers. In doing so it illustrates a reversal of the baseline shift hypothesis that has been proposed for many fisheries (Pauly 1995), whereby generational knowledge is lost and aspirational targets are downwards-adjusted, unintentionally, in response to lived experience. The post-earthquake abundances observed in shallow water provide a window to the past through which baseline shifts can be recognised, and possibilities illustrated that might otherwise remain unknown and unexplored as management objectives.

## 5. Conclusions

The overarching context for this study involves recovery from a natural disaster that illuminated both the nature of an open-access fishery and its potential for recovery and longer-term sustainable use. The key insights gained, which are transferable to other fisheries, include the need to effectively address recreational fishing pressure if relying on daily bag limits, and the utility of temporal closure strategies in the design of solutions that are complementary to protected areas and can support adaptive management by enabling responsive strategies such as flexible closing dates. Their implementation, however, requires greater attention to simple measures such as the registration or licensing of fishers, and the collection of information on recreational catches and effort, all of which could improve the understanding of recreational fisheries but are often overlooked in the management of marine fisheries in New Zealand and elsewhere. These strategic directions also require more meaningful engagement with recreational fishers and more participatory data collection, but are becoming easier to implement thanks to advances in mobile technologies.

Equitable benefit-sharing between commercial and non-commercial sectors requires the reliable implementation of total allowable catches, and can include equity considerations within the recreational sector that arise from patterns in fishing pressure that are enabled, either intentionally or unintentionally, from the choice of management settings. Desirable attributes also include support for periodic stock recovery following unexpected losses from either natural or man-made circumstances. These re-sets present opportunities for re-thinking management objectives, and as exemplified in this study, can offer new insights that include the reversal of baseline shifts that arise from the prevalence of degraded conditions. Taking advantage of the learning available from these events has the potential to improve both current outcomes and future resilience. These principles are generally applicable to many open-access fisheries, and are likely to be particularly valuable where recreational fishing is not constrained through a tag-based system or other directly enforceable limit on total long-term catches.

## Acknowledgements

This research contributes to evaluations of earthquake impacts and recovery status for the Pāua Industry Council (PIC) and Fisheries New Zealand (FNZ), and extends previous work on wider coastal environment recovery funded by the Ministry for Business and Innovation (MBIE) and Ministry for Primary Industries (MPI). We thank Derek Gerber, Zoe Smeele and Jan McKenzie for field assistance with these studies, and acknowledge the considerable work of the many people and organisations that have provided inputs into discussions and decisions on the future of this fishery.

## Footnotes

1 New Zealand Recreational Fishing Council Inc v Sanford [2009] NZSC 54

## References

Abbott, J. K., & Haynie, A. C. (2012). What are we protecting? Fisher behavior and the unintended consequences of spatial closures as a fishery management tool. Ecological Applications, 22(3), 762–777. doi:10.1890/11-1319.1

Arlinghaus, R., Cooke, S. J., & Potts, W. (2013). Towards resilient recreational fisheries on a global scale through improved understanding of fish and fisher behaviour. Fisheries Management and Ecology, 20(2-3), 91–98. doi:10.1111/fme.12027

Arlinghaus, R., Tillner, R., & Bork, M. (2015). Explaining participation rates in recreational fishing across industrialised countries. Fisheries Management and Ecology, 22(1), 45–55. doi:10.1111/fme.12075

Arlinghaus, R., Abbott, J. K., Fenichel, E. P., Carpenter, S. R., Hunt, L. M., Alós, J.,… Manfredo, M. J. (2019). Governing the recreational dimension of global fisheries. Proceedings of the National Academy of Sciences, 116(12), 5209–5213. doi:10.1073/pnas.1902796116

Barbier, E. B., Hacker, S. D., Kennedy, C., Koch, E. W., Stier, A. C., & Silliman, B. R. (2011). The value of estuarine and coastal ecosystem services. Ecological Monographs, 81(2), 169–193. doi:10.1890/10-1510.1

Bastardie, F., Nielsen, J. R., Eigaard, O. R., Fock, H. O., Jonsson, P., & Bartolino, V. (2014). Competition for marine space: modelling the Baltic Sea fisheries and effort displacement under spatial restrictions. ICES Journal of Marine Science, 72(3), 824–840. doi:10.1093/icesjms/fsu215

Beddington, J. R., Kirkwood, G. P., Cochrane, K. L., & Doulman, D. J. (2005). The rising tide of fisheries instruments and the struggle to keep afloat. Philosophical Transactions of the Royal Society B: Biological Sciences, 360(1453), 77–94. doi:doi:10.1098/rstb.2004.1568

Begossi, A. (2014). Ecological, cultural, and economic approaches to managing artisanal fisheries. Environment Development and Sustainability, 16(1), 5–34. doi:10.1007/s10668-013-9471-z

Bohnsack, J. A. (1998). Application of marine reserves to reef fisheries management. Australian Journal of Ecology, 23, 298–304.

Carbines, G. (2000). Kaikoura recreational fishing survey (1998/99). Report prepared for Ministry of Fisheries. 41pp.

Cooke, S. J., & Cowx, I. G. (2006). Contrasting recreational and commercial fishing: Searching for common issues to promote unified conservation of fisheries resources and aquatic environments. Biological Conservation, 128(1), 93–108. doi:10.1016/j.biocon.2005.09.019

Douvere, F. (2008). The importance of marine spatial planning in advancing ecosystem-based sea use management. Marine Policy, 32(5), 762–771. doi:10.1016/j.marpol.2008.03.021

Dyck, A. J., & Sumaila, U. R. (2010). Economic impact of ocean fish populations in the global fishery. Journal of Bioeconomics, 12, 227–243. doi:10.1007/s10818-010-9088-3

Falconer, T. R. L., Gerrity, S., Dunmore, R. A., Crossett, D., Orchard, S., & Schiel, D. R. (2022). Rocky reef impacts of the 2016 Kaikoura earthquake: extended monitoring of nearshore habitats and communities to 5.5 years.Unpublished Progress Report prepared for Fisheries New Zealand project KAI2020-01. Fisheries New Zealand, Wellington. 50pp.

Fisheries New Zealand. (2021). Review of Sustainability Measures for Pāua (PAU 3A & PAU 3B) for 2021/22. Fisheries NZ Discussion Paper No: 2021/13. Wellington: Fisheries New Zealand. 15pp.

Folke, C., Hahn, T., Olsson, P., & Norberg, J. (2005). Adaptive governance of Social-Ecological Systems. Annual Review of Environment and Resources, 30(1), 441–473. doi:10.1146/annurev.energy.30.050504.144511

Folke, C. (2007). Social–ecological systems and adaptive governance of the commons. Ecological Research, 22(1), 14–15. doi:10.1007/s11284-006-0074-0

Fu, D., McKenzie, A., & Marsh, C. (2017). Summary of input data for the 2016 PAU 5D stock assessment. New Zealand Fisheries Assessment Report 2017/32. 79pp.

Gaines, S. D., White, C., Carr, M. H., & Palumbi, S. R. (2010). Designing marine reserve networks for both conservation and fisheries management. Proceedings of the National Academy of Sciences, 107(43), 18286–18293. doi:doi:10.1073/pnas.0906473107

Garcia, S. M., & Charles, A. T. (2008). Fishery systems and linkages: Implications for science and governance. Ocean & Coastal Management, 51(7), 505–527. doi:10.1016/j.ocecoaman.2008.05.001

Gerrity, S., Alestra, T., Fischman, H. S., & Schiel, D. R. (2020). Earthquake effects on abalone habitats and populations in southern New Zealand. Marine Ecology Progress Series, 656, 153–161.

Gerrity, S., & Schiel, D. R. (2022). Fishing effects of wade-able pāua populations along the Kaikoura coast. New Zealand Fisheries Assessment Report 2022/SEA2021-07. Report prepared for Fisheries New Zealand. 37pp.

Government of Western Australian Department of Fisheries. (2021). Abalone resource of Western Australia: Harvest Strategy: 2021-2026: Version 2.0. Government of Western Australia Department of Fisheries, Perth. 46pp. Retrieved 1 October 2022 from https://library.dpird.wa.gov.au/cgi/viewcontent.cgi?article=1308&context=fr_fmp.

Haasnoot, M., Kwakkel, J. H., Walker, W. E., & ter Maat, J. (2013). Dynamic adaptive policy pathways: A method for crafting robust decisions for a deeply uncertain world. Global Environmental Change, 23(2), 485–498. doi:10.1016/j.gloenvcha.2012.12.006

Halpern, B. S., Lester, S. E., & Kellner, J. B. (2010). Spillover from marine reserves and the replenishment of fished stocks. Environmental Conservation, 36(4), 268–276. doi:10.1017/S0376892910000032

Hardin, G. (1968). The Tragedy of the Commons. Science, 162(3859), 1243–1248.

Hart, A. M., Strain, L. W. S., & Brown, J. (2018). Regulation dynamics of exploited and protected populations of Haliotis roei, and their response to a marine heatwave. ICES Journal of Marine Science, 75(6), 1924–1939. doi:10.1093/icesjms/fsy064

Heinemann, A., Wynne-Jones, J., Gray, A., & Hill, L. (2015). National panel survey of marine recreational fishers 2011–12 rationale and methods. New Zealand Fisheries Assessment Report 2015/48. 94pp.

Hinkel, J., Bots, P. W. G., & Schlüter, M. (2014). Enhancing the Ostrom social-ecological system framework through formalization. Ecology and Society, 19(3). doi:10.5751/ES-06475-190351

Holdsworth, J. C. (2022). Harvest estimates from land-based amateur fishers—Kaikōura Marine Area to Marfells Beach. New Zealand Fisheries Assessment Report 2022/40. Report prepared for Fisheries New Zealand. 27pp.

Horta e Costa, B., Batista, M. I., Gonçalves, L., Erzini, K., Caselle, J. E., Cabral, H. N., & >Gonçalves, E. J. (2013). Fishers’ Behaviour in Response to the Implementation of a Marine Protected Area. PLOS ONE, 8(6), e65057. doi:10.1371/journal.pone.0065057

Ihde, T. F., Wilberg, M. J., Loewensteiner, D. A., Secor, D. H., & Miller, T. J. (2011). The increasing importance of marine recreational fishing in the US: Challenges for management. Fisheries Research, 108(2), 268–276. doi:10.1016/j.fishres.2010.12.016

Jamieson, G. S. (1993). Marine invertebrate conservation: Evaluation of fisheries over-exploitation concerns. American Zoologist, 33(6), 551–567. doi:10.1093/icb/33.6.551

Jennings, S. (2000). Patterns and prediction of population recovery in marine reserves. Reviews in Fish Biology and Fisheries, 10(2), 209–231. doi:10.1023/A:1016619102955

Kockel, A., Ban, N. C., Costa, M., & Dearden, P. (2020). Addressing distribution equity in spatial conservation prioritization for small-scale fisheries. PloS one, 15(5), e0233339. doi:10.1371/journal.pone.0233339

Lewis, S. G. (2015). Bags and tags: randomized response technique indicates reductions in illegal recreational fishing of red abalone (*Haliotis rufescens*) in Northern California. Biological Conservation, 189, 72–77. doi:10.1016/j.biocon.2014.09.024

MacKinnon, K., Dudley, N., & Sandwith, T. (2011). Natural solutions: protected areas helping people to cope with climate change. Oryx, 45(4), 461–462. doi:10.1017/S0030605311001608

McCarthy, A., Hepburn, C., Scott, N., Schweikert, K., Turner, R., & Moller, H. (2014). Local people see and care most? Severe depletion of inshore fisheries and its consequences for Māori communities in New Zealand. Aquatic Conservation: Marine and Freshwater Ecosystems, 24(3), 369–390.

McGinnis, M. D., & Ostrom, E. (2014). Social-ecological system framework: initial changes and continuing challenges. Ecology and Society, 19(2). doi:10.5751/ES-06387-190230

McShane, P. E. (1992). Early life history of abalone: a review. In M. J. T. S.A. Shepherd, and S. Guzmán del Próo (Ed.), Abalone of the world: Biology, fisheries and culture (pp. 120–138). Oxford, U.K.: Blackwell Scientific.

Montgomery, M., & Vaughan, M. (2018). Ma Kahana ka ‘Ike: Lessons for community-based fisheries management. Sustainability (Switzerland), 10(10), 3799. doi:10.3390/su10103799

Morales-Nin, B., Cardona-Pons, F., Maynou, F., & Grau, A. M. (2015). How relevant are recreational fisheries? Motivation and activity of resident and tourist anglers in Majorca. Fisheries Research, 164, 45–49. doi:10.1016/j.fishres.2014.10.010

Neubauer, P. (2020). Development and application of a spatial stock assessment model for pāua (Haliotis iris). New Zealand Fisheries Assessment Report 2020/30. Report prepared for Fisheries New Zealand. 42pp.

New Zealand Government. (2021). Kaikōura (Te Tai o Marokura) Marine Management Act 2014. Version as at 28 October 2021. Wellington: New Zealand Government. Available at https://www.legislation.govt.nz/act/public/2014/0059/latest/LMS559230.html.

New Zealand Government. (2022). Fisheries Act 1996. Version as at 1 July 2022. Wellington: New Zealand Government. Available at https://www.legislation.govt.nz/act/public/1996/0088/latest/DLM394192.html.

Orchard, S., Fischman, H. S., Gerrity, S., Alestra, T., Dunmore, R., & Schiel, D. R. (2021). Threshold effects of relative sea-level change in intertidal ecosystems: empirical evidence from earthquake-induced uplift on a rocky coast. GeoHazards, 2(4), 302–320. doi:10.3390/geohazards2040016

Ostrom, E. (1990). Governing the commons: the evolution of institutions for collective action. Cambridge;New York;: Cambridge University Press.

Pauly, D. (1995). Anecdotes and the shifting baseline syndrome of fisheries. Trends in Ecology and Evolution, 10(10), 430. doi:10.1016/s0169-5347(00)89171-5

Pita, C., Villasante, S., & Pascual-Fernández, J. J. (2019). Managing small-scale fisheries under data poor scenarios: lessons from around the world. Marine Policy, 101, 154–157. doi:10.1016/j.marpol.2019.02.008

Radford, Z., Hyder, K., Zarauz, L., Mugerza, E., Ferter, K., Prellezo, R.,… Weltersbach, M. S. (2018). The impact of marine recreational fishing on key fish stocks in European waters. PLOS ONE, 13(9), e0201666. doi:10.1371/journal.pone.0201666

Raymond-Yakoubian, J., Raymond-Yakoubian, B., & Moncrieff, C. (2017). The incorporation of traditional knowledge into Alaska federal fisheries management. Marine Policy, 78, 132–142. doi:10.1016/j.marpol.2016.12.024

Schiel, D. R., & Breen, P. A. (1991). Population structure, ageing and fishing mortality of the New Zealand abalone *Haliotis iris*. Fishery Bulletin, 89, 681–691.

Schiel, D. R. (1992). *The paua (abalone) fishery of New Zealand*. In: Abalone of the world: Biology,fisheries and culture. Shepherd, S.A.; Tegner, M.J.; Guzman del Proo, S. (eds.) pp. 427–437. Oxford: Blackwell Scientific.

Schiel, D. R., Andrew, N. L., & Foster, M. S. (1995). The structure of subtidal algal and invertebrate assemblages at the Chatham Islands, New Zealand. Marine Biology, 123(2), 355–367. doi:10.1007/BF00353627

Schiel, D. R., Alestra, T., Gerrity, S., Orchard, S., Dunmore, R., Pirker, J.,… Thomsen, M. (2019). The Kaikōura earthquake in southern New Zealand: loss of connectivity of marine communities and the necessity of a cross-ecosystem perspective. Aquatic Conservation: Marine and Freshwater Ecosystems, 29(9), 1520–1534. doi:10.1002/aqc.3122

Schiel, D. R., Gerrity, S., Orchard, S., Alestra, T., Dunmore, R. A., Falconer, T.,… Tait, L. W. (2021). Cataclysmic disturbances to an intertidal ecosystem: loss of ecological infrastructure slows recovery of biogenic habitats and diversity. Frontiers in Ecology and Evolution, 9(827). doi:10.3389/fevo.2021.767548

Strain, E. M. A., Alexander, K. A., Kienker, S., Morris, R., Jarvis, R., Coleman, R.,… Bishop, M. J. (2019). Urban blue: A global analysis of the factors shaping people’s perceptions of the marine environment and ecological engineering in harbours. Science of the Total Environment, 658, 1293–1305. doi:10.1016/j.scitotenv.2018.12.285

Strain, L., Brown, J., & Jones, R. (2021). West Coast Roe’s Abalone Resource Status Report. In: Status Reports of the Fisheries and Aquatic Resources of Western Australia 2019/20: State of the Fisheries (pp. 37–42): Department of Primary Industry and Regional Development, Western Australia.

Thomsen, M. S., Mondardini, L., Thoral, F., Gerber, D., Montie, S., South, P. M.,… Schiel, D. R. (2021). Cascading impacts of earthquakes and extreme heatwaves have destroyed populations of an iconic marine foundation species. Diversity and Distributions, 27(12), 2369–2383. doi:10.1111/ddi.13407

Watson, J. E. M., Dudley, N., Segan, D. B., & Hockings, M. (2014). The performance and potential of protected areas. Nature, 515(7525), 67–73. doi:10.1038/nature13947

Wilson, N. H. F., & Schiel, D. R. (1995). Reproduction of two species of abalone (*Haliotis iris* and *H. australis*) in southern New Zealand. Marine and Freshwater Research, 46(3), 629–637. doi:10.1071/MF9950629

Yang, G., Ge, Y., Xue, H., Yang, W., Shi, Y., Peng, C.,… Chang, J. (2015). Using ecosystem service bundles to detect trade-offs and synergies across urban–rural complexes. Landscape and Urban Planning, 136(0), 110–121. doi:10.1016/j.landurbplan.2014.12.006

