## Supplementary material for "Re-thinking recreational fishing – how a natural disaster presents insights and opportunities for achieving sustainability and equity objectives": Table S1

**Table S1.** Fisher numbers as observed during coast-wide field surveys completed between Oaro and Waipapa Bay in the 2021 pāua fishing season on the Kaikōura coast.

| **Date** | **Period** | **Observed fishers** | **Visitation coefficient†** | **Estimated fishers** | **Estimated catch*** | **Start_time** | **Low_tide** | **Tide_height** | **Wind_dir** | **Wind_str** | **Precip** | **Cloud** | **MaxTemp** | **Swell_m** | | **Swell_dir** |
| --- | --- | --- | --- | --- | --- | --- | --- | --- | --- | --- | --- | --- | --- | --- | --- | --- |
| 08/12/2021 | weekday | 56 | 1.5 | 84 | 420 | 1430 | 1511 | 0.3 | E | L | nil | 2 | 19 | 1 | | NE |
| 09/12/2021 | weekday | 40 | 1.5 | 60 | 300 | 1500 | 1610 | 0.3 | SW | L | nil | 2 | 18 | 1.5 | | NE |
| 10/12/2021 | weekday | 68 | 1.5 | 102 | 510 | 1645 | 1707 | 0.3 | NE | L | nil | 2 | 19 | 1 | | NE |
| 18/12/2021 | weekend | 57 | 1.5 | 86 | 428 | 1000 | 1127 | 0.6 | NE | L | nil | 0 | 17 | 0.5 | | NE |
| 20/12/2021 | weekday | 89 | 1.5 | 134 | 668 | 1130 | 1253 | 0.6 | E | L | drizzle | 4 | 20 | 0.5 | | NE |
| 21/12/2021 | weekday | 122 | 1.5 | 183 | 915 | 1200 | 1336 | 0.6 | SE | L | drizzle | 4 | 18 | 0.5 | | NE |
| 22/12/2021 | weekday | 140 | 1.5 | 210 | 1050 | 1230 | 1418 | 0.6 | E | M | rain | 4 | 18 | 1.5 | | NE |
| 23/12/2021 | weekday | 116 | 1.5 | 174 | 870 | 1330 | 1500 | 0.6 | S | M | nil | 3 | 20 | 1 | | NE |
| 27/12/2021 | holiday | 305 | 2.5 | 763 | 3813 | 1600 | 1757 | 0.5 | S | M | nil | 2 | 22 | 1 | | SE |
| 02/01/2022 | holiday | 497 | 2.5 | 1243 | 6213 | 0900 | 1059 | 0.3 | SW | L | nil | 1 | 21 | 0.5 | | NE |
| 10/01/2022 | holiday | 199 | 2.5 | 498 | 2488 | 1700 | 1625 | 0.5 | N | L | nil | 1 | 24 | 0.5 | | NE |
| 11/10/2022 | holiday | 138 | 2.5 | 345 | 1725 | 1730 | 1914 | 0.5 | NE | L | nil | 1 | 25 | 0.5 | | NE |
| 26/01/2022 | weekday | 54 | 1.5 | 81 | 405 | 1700 | 1818 | 0.5 | E | L | nil | 2 | 19 | 1 | | E |
| 27/01/2022 | weekday | 11 | 1.5 | 17 | 83 | 1730 | 1909 | 0.4 | SW | M | nil | 2 | 19 | 1.5 | | S |
| 28/01/2022 | weekday | 27 | 1.5 | 41 | 203 | 1830 | 2004 | 0.4 | NE | L | nil | 2 | 22 | 1.5 | | SE |
| 29/01/2022 | weekend | 152 | 2.0 | 304 | 1520 | 0730 | 0847 | 0.4 | NW | L | nil | 1 | 22 | 1 | | E |
| 18/02/2022 | weekday | 77 | 1.5 | 116 | 578 | 1200 | 1307 | 0.5 | N | L | nil | 1 | 20 | 0.5 | | E |
| 19/02/2022 | weekend | 394 | 2.0 | 788 | 3940 | 1300 | 1352 | 0.5 | NW | L | nil | 2 | 20 | 0.5 | | E |
| 20/02/2022 | weekend | 94 | 2.0 | 188 | 940 | 1345 | 1438 | 0.5 | W | L | nil | 1 | 16 | 0.5 | | E |
| **TOTALS** |  |  |  |  | **27065** | **pāua** |  |  |  |  |  |  |  |  |  |  |
|  |  |  |  |  | **9.02** | **tonne** |  |  |  |  |  |  |  |  |  |  |

† reflects the difference between the fisher numbers observed during a short (c. 5 minute) period at each location and total daily fisher numbers as assessed using time series data from representative sites

* based on an assumed catch-per-day (CPD) of 5 pāua per person
