## Supplementary material for "Re-thinking recreational fishing – how a natural disaster presents insights and opportunities for achieving sustainability and equity objectives": Table S2

**Table S2**. Uncertainty estimates for seasonal fishing pressure model

|  | Daily fishing pressure (number of fishers) | | | | | |
| --- | --- | --- | --- | --- | --- | --- |
| **Period** | **Best estimate** | | **Lower bound** | | **Upper bound** | |
|  | **Visitation coefficient** | **Mean ± SE** | **Visitation coefficient** | **Mean ± SE** | **Visitation coefficient** | **Mean ± SE** |
| holiday | 2.5 | 712 ± 176 | 2.0 | 569 ± 157 | 3.0 | 854 ± 236 |
| weekend | 2.0 | 341 ± 155 | 1.5 | 254 ± 118 | 2.5 | 429 ± 193 |
| weekday | 1.5 | 109 ± 19 | 1.0 | 73 ± 12 | 2.0 | 146 ± 25 |
