## Supplementary material for "Re-thinking recreational fishing – how a natural disaster presents insights and opportunities for achieving sustainability and equity objectives": Table S3

**Table 3**. Daily weather observations for Kaikōura AWS Station over the 2021 recreational pāua fishing season. Poor weather days included in the seasonal catch model are in shown bold. Source: National Climate Database.

| **Date** | **Temp_min (^o^C)** | **Temp_max (^o^C)** | **Rainfall  (mm)** | **MaxGust (m/s)** |
| --- | --- | --- | --- | --- |
| 20211201 | 11.8 | 16.6 | 0 | 9.3 |
| 20211202 | 12.5 | 16.9 | 0.2 | 11.8 |
| 20211203 | 13.6 | 16.2 | 0.6 | 5.7 |
| 20211204 | 16 | 22.4 | 0.2 | 7.7 |
| 20211205 | 18 | 25.8 | 0 | 20.1 |
| 20211206 | 9.9 | 27.7 | 5.4 | 19 |
| **20211207** | **9.5** | **12.4** | **2.6** | **6.2** |
| 20211208 | 11.5 | 15.1 | 0 | 6.7 |
| 20211209 | 13.8 | 19.2 | 0 | 11.3 |
| 20211210 | 14 | 20.1 | 0 | 14.4 |
| 20211211 | 14.1 | 17.6 | 2.4 | 18.5 |
| **20211212** | **9.6** | **-** | **28.2** | **13.4** |
| **20211213** | **10.1** | **13.9** | **0.2** | **9.8** |
| 20211214 | 13.7 | 17.3 | 0 | 11.8 |
| 20211215 | 12.3 | 14.4 | 67.2 | 13.9 |
| 20211216 | 13.1 | 16.8 | 55.8 | 18.5 |
| 20211217 | 12.7 | 19.1 | 0.2 | 16.5 |
| 20211218 | 10 | 15.3 | 0 | 10.8 |
| 20211219 | 11.8 | 19 | 0 | 9.3 |
| 20211220 | 15.4 | 23.8 | 0 | 12.9 |
| 20211221 | 17.8 | 26.4 | 0 | 14.4 |
| 20211222 | 15.7 | 23.4 | 0 | 14.9 |
| 20211223 | 15.1 | 25.9 | 0 | 17.5 |
| 20211224 | 14.7 | 18.8 | 0 | 14.9 |
| 20211225 | 13.1 | 19.3 | 0 | 15.4 |
| 20211226 | 11.2 | 17.1 | 0.8 | 7.2 |
| 20211227 | 16.3 | 22.2 | 0 | 11.8 |
| 20211228 | 14.4 | 25.3 | 0.4 | 15.4 |
| 20211229 | 11.2 | 16.5 | 28.6 | 11.8 |
| 20211230 | 11.2 | 16.2 | 0.8 | 11.3 |
| 20211231 | 12.7 | 20.5 | 6.8 | 11.8 |
| 20220101 | 13.9 | 20.9 | 0 | 6.7 |
| 20220102 | 14.2 | 24.4 | 0 | 5.7 |
| 20220103 | 17 | 23 | 0 | 7.2 |
| 20220104 | 15.1 | 28.5 | 0 | 6.2 |
| 20220105 | 15.7 | 19 | 0 | 9.8 |
| 20220106 | 11.9 | 20.4 | 0.6 | 19.6 |
| 20220107 | 11 | 14.6 | 1.6 | 14.9 |
| 20220108 | 11 | 16.6 | 0 | 8.8 |
| 20220109 | 15.1 | 20.5 | 0.2 | 7.7 |
| 20220110 | 17.7 | 25.5 | 0 | 7.2 |
| 20220111 | 16 | 23.9 | 0 | 16 |
| 20220112 | 15.5 | 26.1 | 0 | 19.6 |
| 20220113 | 10.5 | 18.7 | 4.4 | 15.4 |
| 20220114 | 10.4 | 16.3 | 0.6 | 6.7 |
| 20220115 | 12 | 17.5 | 0 | 7.7 |
| 20220116 | 12.1 | 19.6 | 0 | 12.4 |
| 20220117 | 13.2 | 20.6 | 0 | 9.3 |
| 20220118 | 13.9 | 21.9 | 0 | 9.3 |
| 20220119 | 13.7 | 21.5 | 0 | 14.9 |
| 20220120 | 10.6 | 26.5 | 18.4 | 20.1 |
| 20220121 | 10.6 | 16.8 | 0.2 | 10.8 |
| 20220122 | 12.8 | 19 | 0 | 9.8 |
| 20220123 | 15.3 | 21.7 | 0.4 | 8.8 |
| 20220124 | 13.3 | 16.8 | 15.8 | 12.4 |
| 20220125 | 14.4 | 19.2 | 0.8 | 7.2 |
| 20220126 | 15.6 | 19.1 | 0 | 31.4 |
| 20220127 | 10.2 | 20.8 | 6.4 | 30.9 |
| 20220128 | 11.8 | 16.3 | 0 | 11.8 |
| 20220129 | 11.2 | 18.4 | 0 | 8.8 |
| 20220130 | 14.9 | 22.6 | 0 | 10.3 |
| 20220131 | 14.6 | 22.3 | 0 | 12.4 |
| 20220201 | 15.1 | 19.6 | 0 | 6.2 |
| 20220202 | 17.1 | 24.4 | 0 | 6.7 |
| 20220203 | 20.7 | 26.2 | 1.4 | 12.4 |
| 20220204 | 14.6 | 27.6 | 1.6 | 12.9 |
| 20220205 | 12.9 | 17.8 | 3.8 | 12.4 |
| 20220206 | 11.6 | 14.6 | 70 | 10.3 |
| **20220207** | **10.7** | **13.8** | **3.8** | **7.2** |
| 20220208 | 12 | 17.6 | 4 | 5.1 |
| 20220209 | 13.3 | 16.8 | 0.8 | 5.7 |
| 20220210 | 15.1 | 19.7 | 19.8 | 6.7 |
| 20220211 | 16.1 | 23 | 0 | 8.8 |
| 20220212 | 15.2 | 18.9 | 0 | 13.9 |
| 20220213 | 9.2 | 16.5 | 76 | 23.2 |
| **20220214** | **9.1** | **12.1** | **32.6** | **16.5** |
| 20220215 | 11.5 | 17.1 | 0 | 8.8 |
| 20220216 | 11.3 | 19 | 0.2 | 18 |
| 20220217 | 12.9 | 19.7 | 0 | 8.2 |
| 20220218 | 12.7 | 19.2 | 0 | 4.6 |
| 20220219 | 16.1 | 23.3 | 0.2 | 23.2 |
| 20220220 | 10.6 | 25.3 | 23.2 | 12.4 |
| 20220221 | 11.7 | 17.1 | 0 | 12.4 |
| 20220222 | 14.2 | 19 | 0 | 26.8 |
| 20220223 | 13.6 | 26.3 | 0 | 19.6 |
| 20220224 | 12.2 | 16.3 | 0.2 | 7.2 |
| 20220225 | 10 | 17.1 | 0 | 9.8 |
| 20220226 | 11.1 | 18.8 | 0 | 20.1 |
| 20220227 | 10.3 | 18.2 | 0.2 | 12.4 |
| **20220228** | **8.9** | **14.1** | **0** | **7.7** |
