## Supplementary material for "Re-thinking recreational fishing – how a natural disaster presents insights and opportunities for achieving sustainability and equity objectives": Table S4

**Table S4** Seasonal catch scenarios for combinations of catch-per-day (CPD) limits, No Fishing Day (NFD) and seasonal timing adjustments relative to a base-case that reflects the pattern of fishing pressure observed in the 2021-2022 open season on the Kaikōura coast. The equivalent 2021-2022 season CPD limit is shown in grey.

(a) No Fishing Days = nil [shortened season + CPD combinations only]

| **No Fishing days** | **Length of season (weeks)** | **Closing  date** | **Seasonal catch (tonnes)** | | | | | |
| --- | --- | --- | --- | --- | --- | --- | --- | --- |
|  |  |  | **CPD1** | **CPD2** | **CPD3** | **CPD4** | **CPD5** | **CPD6** |
| nil | 13 | 28 February | 9.0 | 18.0 | 27.0 | 36.0 | 45.0 | 54.0 |
| nil | 12 | 20 February | 8.6 | 17.2 | 25.7 | 34.3 | 42.9 | 51.5 |
| nil | 11 | 13 February | 8.2 | 16.4 | 24.6 | 32.8 | 41.0 | 49.3 |
| nil | 10 | 6 February | 7.8 | 15.7 | 23.5 | 31.3 | 39.2 | 47.0 |
| nil | 9 | 30 January | 7.5 | 14.9 | 22.4 | 29.9 | 37.3 | 44.8 |

(b) No Fishing Day = Mon

| **No Fishing days** | **Length of season (weeks)** | **Closing  date** | **Seasonal catch (tonnes)** | | | | | |
| --- | --- | --- | --- | --- | --- | --- | --- | --- |
|  |  |  | **CPD1** | **CPD2** | **CPD3** | **CPD4** | **CPD5** | **CPD6** |
| Mon | 13 | 28 February | 8.1 | 16.1 | 24.2 | 32.2 | 40.3 | 48.4 |
| Mon | 12 | 20 February | 7.6 | 15.2 | 22.7 | 30.3 | 37.9 | 45.5 |
| Mon | 11 | 13 February | 7.2 | 14.4 | 21.6 | 28.8 | 36.0 | 43.2 |
| Mon | 10 | 6 February | 6.8 | 13.7 | 20.5 | 27.3 | 34.1 | 41.0 |
| Mon | 9 | 30 January | 6.5 | 12.9 | 19.4 | 25.8 | 32.3 | 38.7 |

(c) No Fishing Days = Sun + shortened season scenarios

| **No Fishing days** | **Length of season (weeks)** | **Closing  date** | **Seasonal catch (tonnes)** | | | | | |
| --- | --- | --- | --- | --- | --- | --- | --- | --- |
|  |  |  | **CPD1** | **CPD2** | **CPD3** | **CPD4** | **CPD5** | **CPD6** |
| Sun | 13 | 28 February | 7.2 | 14.5 | 21.7 | 29.0 | 36.2 | 43.5 |
| Sun | 12 | 20 February | 6.8 | 13.7 | 20.5 | 27.3 | 34.2 | 41.0 |
| Sun | 11 | 13 February | 6.6 | 13.2 | 19.7 | 26.3 | 32.9 | 39.5 |
| Sun | 10 | 6 February | 6.3 | 12.6 | 19.0 | 25.3 | 31.6 | 37.9 |
| Sun | 9 | 30 January | 6.1 | 12.1 | 18.2 | 24.2 | 30.3 | 36.4 |

(d) Season length scenarios and two No Fishing Days (Sunday+Monday)

| **No Fishing days** | **Length of season (weeks)** | **Closing  date** | **Seasonal catch (tonnes)** | | | | | |
| --- | --- | --- | --- | --- | --- | --- | --- | --- |
|  |  |  | **CPD1** | **CPD2** | **CPD3** | **CPD4** | **CPD5** | **CPD6** |
| Sun+Mon | 13 | 28 February | 6.2 | 12.4 | 18.6 | 24.8 | 31.0 | 37.2 |
| Sun+Mon | 12 | 20 February | 5.8 | 11.7 | 17.5 | 23.3 | 29.1 | 35.0 |
| Sun+Mon | 11 | 13 February | 5.6 | 11.1 | 16.7 | 22.3 | 27.9 | 33.4 |
| Sun+Mon | 10 | 6 February | 5.3 | 10.6 | 15.9 | 21.2 | 26.6 | 31.9 |
| Sun+Mon | 9 | 30 January | 5.1 | 10.1 | 15.2 | 20.2 | 25.3 | 30.3 |

(e) Season length scenarios with seasonal timing to avoid the summer holiday period

| **No Fishing days** | **Length of season (weeks)** | **Open  season** | **Seasonal catch (tonnes)** | | | | | |
| --- | --- | --- | --- | --- | --- | --- | --- | --- |
|  |  |  | **CPD1** | **CPD2** | **CPD3** | **CPD4** | **CPD5** | **CPD6** |
| nil | 13 | Timed to avoid holiday period (late December to end January) | 5.0 | 9.9 | 14.9 | 19.9 | 24.8 | 29.8 |
| nil | 12 |  | 4.6 | 9.2 | 13.8 | 18.3 | 22.9 | 27.5 |
| nil | 11 |  | 4.2 | 8.4 | 12.6 | 16.8 | 21.0 | 25.2 |
| nil | 10 |  | 3.8 | 7.6 | 11.5 | 15.3 | 19.1 | 22.9 |
| nil | 9 |  | 3.4 | 6.9 | 10.3 | 13.8 | 17.2 | 20.6 |

† relative to the 2021-2022 season which opened 1 December 2021 and closed 28 February 2022.
